# HLA*PRG:LA – HLA typing from linearly projected graph alignments

**DOI:** 10.1101/453555

**Authors:** Alexander T Dilthey, Alexander J Mentzer, Raphael Carapito, Clare Cutland, Nezih Cereb, Shabir A. Madhi, Arang Rhie, Sergey Koren, Seiamak Bahram, Gil McVean, Adam M Phillippy

## Abstract

**Summary:** HLA*PRG:LA implements a new graph alignment model for HLA type inference, based on the projection of linear alignments onto a variation graph. It enables accurate HLA type inference from whole-genome (99% accuracy) and whole-exome (93% accuracy) Illumina data; from long-read Oxford Nanopore and Pacific Biosciences data (98% accuracy for whole-genome and targeted data); and from genome assemblies. Computational requirements for a typical sample vary between 0.7 and 14 CPU hours per sample.

**Availability and Implementation:** HLA*PRG:LA is implemented in C++ and Perl and freely available from https://github.com/DiltheyLab/HLA-PRG-LA (GPL v3).

**Supplementary information:** Supplementary data are available online.

## Introduction

Genetic variation at the Human Leukocyte Antigen (HLA) loci is associated with many important phenotypes and biological conditions, including autoimmune and infectious disease risk, transplant rejection, and the repertoire of immune-presented peptides [1]. With the growing availability of whole-exome and whole-genome sequencing data, the ability to accurately determine the allelic state of the HLA genes (“HLA typing”) from these data types is becoming increasingly important. The accuracy of standard sequencing data analysis methods in the HLA region is limited by hyperpolymorphism and reference divergence [2]. Specialized HLA analysis methods, however, have been developed [3-8]; these typically rely on variation-aware alignment approaches, e.g. genome graphs or collections of linear reference sequences. When applied to high-coverage whole-genome sequencing (WGS) data, the accuracy of these methods can approach that of gold-standard HLA typing assays like sequence-based typing (SBT). HLA type inference from exome or low-coverage WGS data, however, remains more challenging, and most tools do not support long-read data.

We present HLA*PRG:LA (“linear alignments”), a graph-based method with high accuracy on exome and low-coverage WGS data, full support for assembled and unassembled long-read data, and a new projection-based approach to graph alignment. Briefly, the alignment process starts with the identification of linear alignments between the input reads and the reference haplotypes that the graph was constructed from; these are projected onto the graph and optimized in a stepwise process. The intuition behind this is that the (projected) original linear alignment will often be a close approximation to the best graph alignment, except for reads that switch between divergent reference haplotypes. In HLA*PRG:LA, these are identified heuristically and trigger a switch into full graph alignment mode. Its hybrid approach enables HLA*PRG:LA to often avoid the - potentially costly - full graph alignment, to leverage the performance of highly optimized linear alignment algorithms, and to carry out graph alignment for both short and long reads in a unified framework.

## Materials and methods

### Read-to-graph alignment and inference

HLA*PRG:LA employs a Population Reference Graph (PRG; [11]) of 13491 sequences, representing the eight GRCh38 [9] MHC haplotypes and the IMGT exonic and genomic sequences [10]. Linear alignments are obtained by aligning input reads against a modified reference genome (GRCh38 plus the eight MHC haplotypes and IMGT genomic sequences) with BWA-MEM [11]. Alignments from regions covered by the PRG are projected onto the graph and undergo a three-stage optimization process.

First, inspection: alignments are split in areas that contain imbalanced gaps (i.e. unequal numbers of insertions and deletions), which can indicate missed haplotype switch points and problems with the global homology structure of the alignment (i.e. which base of the read aligns to which level of the PRG). Second, polishing: balanced insertion-deletion-structures are folded into the existing alignments, and the highest-scoring graph traversal compatible with the global homology structure of the alignment is found for the aligned sequence using dynamic programming. This step integrates small variants that were not part of the linear sequences used for the initial mapping. Third, extension: unaligned bases are integrated by extending the alignment in full graph alignment mode [3].

HLA type inference follows the likelihood model of HLA*PRG [3] and is based on maximizing P(aligned reads|a_1_, a_2_) over all pairs (a_1_, a_2_) of possible alleles at each locus.

For long-read input data, the inspection and extension steps are skipped, and the likelihood model is adjusted for higher error rates.

Alignment and inference algorithms are illustrated in Supplementary Figure S1.

### Assembly typing

HLA typing of assemblies is based on a projection of the GRCh38 MHC reference haplotype annotations onto the assembly. Briefly, MHC-overlapping contigs are identified with nucmer [12]. For each identified contig and each reference haplotype [9], a semi-global alignment is computed heuristically (see below); the annotations of the haplotype underlying the highest-scoring alignment are projected onto the contig; HLA gene and exon sequences are extracted based on the projected coordinates. HLA typing is carried out by IMGT database matching (minimum edit distance). The global alignment heuristic uses BWA-MEM for the identification of local alignments (“diagonals”) between the input sequences; dynamic programming is used to identify the highest-scoring traversal of the global alignment matrix limited to the identified diagonals connected with horizontal or vertical jumps. This approach accounts for different MHC haplotype structures and ensures that each gene is used only once per contig.

## Results

We carry out three experiments (Table 1) to assess the performance of HLA*PRG:LA.

**Table 1:** Summary validation results

| Cohort | Locus | N | HLA*PRG |  | Kourami |  | xHLA |  | HLA*PRG:LA |  |
| --- | --- | --- | --- | --- | --- | --- | --- | --- | --- | --- |
|  |  |  | Call Rate | Accuracy | Call Rate | Accuracy | Call Rate | Accuracy | Call Rate | Accuracy |
| Illumina WGS | A | 28 | 1.00 | 1.00 | 1.00 | 0.96 | 1.00 | 1.00 | 1.00 | 1.00 |
|  | B | 28 | 1.00 | 1.00 | 1.00 | 1.00 | 1.00 | 1.00 | 1.00 | 1.00 |
|  | C | 28 | 1.00 | 1.00 | 1.00 | 1.00 | 1.00 | 1.00 | 1.00 | 1.00 |
|  | DQA1 | 18 | 1.00 | 1.00 | 1.00 | 1.00 | 0.00 | NA | 1.00 | 1.00 |
|  | DQB1 | 28 | 1.00 | 1.00 | 1.00 | 1.00 | 1.00 | 1.00 | 1.00 | 1.00 |
|  | DRB1 | 28 | 1.00 | 0.96 | 1.00 | 1.00 | 1.00 | 1.00 | 1.00 | 0.96 |
| Illumina Exome | A | 58 | 1.00 | 0.86 | 0.76 | 0.91 | 1.00 | 0.98 | 1.00 | 0.93 |
|  | B | 48 | 1.00 | 0.85 | 0.63 | 0.87 | 1.00 | 0.88 | 1.00 | 0.92 |
|  | C | 56 | 1.00 | 0.79 | 0.61 | 0.97 | 1.00 | 0.96 | 1.00 | 0.88 |
|  | DQA1 | 58 | 1.00 | 0.95 | 1.00 | 0.97 | 0.00 | NA | 1.00 | 0.97 |
|  | DQB1 | 58 | 1.00 | 0.98 | 0.97 | 0.96 | 1.00 | 0.95 | 1.00 | 0.98 |
|  | DRB1 | 58 | 1.00 | 0.91 | 1.00 | 0.97 | 1.00 | 0.98 | 1.00 | 0.91 |
| Long-read WGS | A | 6 | NA |  | NA |  | NA |  | 1.00 | 1.00 |
|  | B | 6 |  |  |  |  |  |  | 1.00 | 1.00 |
|  | C | 6 |  |  |  |  |  |  | 1.00 | 1.00 |
|  | DQA1 | 6 |  |  |  |  |  |  | 1.00 | 1.00 |
|  | DQB1 | 6 |  |  |  |  |  |  | 1.00 | 1.00 |
|  | DRB1 | 6 |  |  |  |  |  |  | 1.00 | 0.83 |
| Long-read Targeted | A | 360 | NA |  | NA |  | NA |  | 1.00 | 1.00 |
|  | B | 362 |  |  |  |  |  |  | 1.00 | 1.00 |
|  | C | 362 |  |  |  |  |  |  | 1.00 | 1.00 |
|  | DQA1 | 318 |  |  |  |  |  |  | 1.00 | 0.97 |
|  | DQB1 | 362 |  |  |  |  |  |  | 1.00 | 0.97 |
|  | DRB1 | 280 |  |  |  |  |  |  | 1.00 | 0.95 |

On high-coverage Illumina WGS data [13, 14], HLA*PRG:LA achieves a performance of 99.4% averaged over the six classical HLA genes. This is identical to the performance of Kourami [8] and HLA*PRG. xHLA [7] achieves an accuracy of 100%, but does not produce calls for *HLA-DQA1*. The performance of HLA*PRG:LA under reduced coverage (15X) is relatively stable (90%) and comparable to that of xHLA (91%), where xHLA has higher performance on HLA class I (95% compared to 87%), and HLA*PRG:LA has higher performance on HLA class II (94% compared to 84%). Kourami is less stable under reduced coverage (average accuracy 69%).

On whole-exome Illumina sequencing data [15], HLA*PRG:LA outperforms HLA*PRG (93% accuracy compared to 89%) and Kourami at matched call rates (95.3% at 88% call rate compared to 94.6% accuracy at 83% call rate). The overall accuracy of xHLA is 95%, with *HLA-DQA1* remaining uncalled.

On whole-genome and targeted PacBio and Nanopore sequencing data [16-18], HLA*PRG:LA achieves an average accuracy of 98%. Of note, the majority (2034 alleles) of the long-read validation data consists of highly diverse South African samples. HLA*PRG, Kourami and xHLA do not support long-read data.

Complete validation results by cohort, a description of the validation cohorts and accessions, and truth HLA types are given in Supplementary Table S1, Supplementary Note S1 and Supplementary Table S2. Computational requirements depend on sample type and coverage (0.65 CPU hours on average for exome samples; between 2.9 and 14 CPU hours for a typical WGS sample; Supplementary Table S3).

The assembly typing component was successfully applied to assess base-level and diploid MHC assembly and to detect phasing and haplotype errors in two recent *de novo* assembly projects [16, 19].

## Conclusion

In summary, HLA*PRG:LA improves upon the accuracy of its predecessor HLA*PRG, while being 3-10 times faster and extending HLA typing functionality to long reads and assemblies. Generalization of projection-based graph alignment beyond the HLA is a topic for future research.

## Acknowledgements

We thank Tina Graves Lindsay and Washington University in St. Louis for public release of the PacBio NA12878 data. We would also like to thank Adrian V Hill, Manjinder Sandhu and others in his group (Cristina Pomilla) and the study participants and the clinic, laboratory, statistical and support staff at RMPRU for providing the South African data available for this study and for their helpful comments during manuscript preparation. We are grateful to all staff at Histogenetics for their involvement in PacBio and Miseq exon-targeted SBT of the South African samples.

## Supplementary Figure Legends

### Supplementary Figure S1

Illustration of the HLA*PRG:LA inference process. (A) The set of input sequences used to construct the PRG. Note that there is a global, multiple-sequence-alignment-like homology structure between the input sequences and that there are shorter and longer input sequences, corresponding to exonic and genomic / regional input haplotypes. (B) A PRG constructed from the input panel, using a recombination model that collapses all homologous-identical characters. (C) The subset of input sequences that will be used for the linear mapping process. Note that these sequences are identical to the long sequences from panel A, with all gaps removed. (D) An example sequencing read. (E) An alignment between the sequencing read and the first haplotype used for linear mapping. Mismatches and gaps are highlighted in red. (F) The graph projection of the linear read-to-haplotype alignment. The graph traversal path corresponding to the alignment is highlighted in green. (G) The read-to-graph alignment, post-inspection. The inspection step heuristically identifies potential recombination points missed by the linear sequence alignment by examining local alignment structure. Here, two consecutive gaps are interpreted as evidence of a potential recombination point, and the last two bases of the read are removed from the alignment. Unaligned bases of the read are highlighted in gray. Note a corresponding reduction of the graph traversal path. (H) The inspected read-to-graph alignment after polishing. The polishing step scans for local improvements of the sequence-to-graph alignment within the existing homology structure of the alignment. (I) The read-to-graph alignment, post-extension. If there are any unaligned read bases, the alignment is extended in full graph-alignment mode. That is, the polished alignment is used as a seed for the graph alignment extension step. (K) HLA type inference is based on the full set of reads overlapping with the typing-relevant exons.

